# Nuclear morphogenesis: forming a heterogeneous nucleus during embryogenesis

**DOI:** 10.1101/2021.10.14.464356

**Authors:** Albert Tsai, Justin Crocker

**Affiliations:** European Molecular Biology Laboratory, Heidelberg, Germany

## Abstract

An embryo experiences progressively complex spatial and temporal patterns of gene expression that guide the morphogenesis of its body plan as it matures. Using super-resolution fluorescence microscopy in *Drosophila melanogaster* embryos, we observed a similar increase in complexity in the nucleus: the spatial distributions of transcription factors became increasingly heterogeneous as the embryo matured. We also observed a similar trend in chromatin conformation with the establishment of specific histone modification patterns. However, transcription sites of specific genes had distinct local preferences for histone marks separate from the average nuclear trend, depending on the time and location of their expression. These results suggest that reconfiguring the nuclear environment is an integral part of embryogenesis and that the physical organization of the nucleus is a key element in developmental gene regulation.

**Summary statement:** We observed spatial rearrangements in the nucleus during embryo development, progressively forming a heterogeneous nuclear environment, paralleling the increasing complexity of the embryo body as morphogenesis progresses.

## Introduction

During embryogenesis, transcriptional regulation controls the expression of genes that ultimately leads to the patterning of the animal (Long et al., 2016; Mallo and Alonso, 2013; Reiter et al., 2017; Spitz and Furlong, 2012). This regulatory process involves interactions between transcription factors, chromatin regulators, and genome topology. Large-scale genome-wide assays have revealed a diversity of functional regulatory elements and the factors that bind them (Consortium, 2012; Johnson et al., 2007; Kundaje et al., 2015; Stunnenberg et al., 2016; Thurman et al., 2012). Notably, these assays have detected wide-ranging reorginizaitons within the nucleus during cellular differentiation, where longer range interactions may bring together chromatin regions and genes into Topologically Associated Domains (TADs). These observations suggest that cellular differentiation involves transforming a nucleus from a uniform compartment into a heterogeneous and partitioned space. However, these aggregate maps represent signals averaged over populations of cells and therefore mask cellular and regulatory heterogeneity. There is now recognition that cell-to-cell variation of transcription factors, chromatin modifications, and interactions between regulatory elements are critical for animal development (Furlong and Levine, 2018; Shema et al., 2019; Tanay and Regev, 2017). However, a significant challenge in the field has been to pair single-cell and gene techniques with complex tissues and devolpmental systems.

Improvements in imaging technology—including dyes, probes, resolution, and throughput—are advancing capabilities for measuring regulatory interactions in developmental systems at the single cell and locus level (Chen et al., 2018; Garcia et al., 2013; Lim et al., 2018; Mir et al., 2017, 2018; Tsai et al., 2017). These techniques can spatially resolve genome regulation at high resolution (Bintu et al., 2018; Mateo et al., 2019; Shema et al., 2019), opening the door to direct observations of the local transcriptional environments around individual gene loci in specific cells. We have previously found that the distributions of transcription factors are heterogeneous in nuclei of *Drosophila* embryos in a late stage of development and that genes such as *shavenbaby* (*svb*) reside in regions enriched for transcription factors when they are transcriptionally active (Tsai et al., 2017).

However, the origins and functional properties of these transcriptional microenvironments during development remain unclear. When do transcriptional microenvironment form in the nucleus? How are these microenvironments affecting transcriptional regulation? In this survey, we imaged the distributions of transcription factors and histone modifications in *Drosophila melanogaster* embryos at different stages of development. Additionally, we imaged histone modifications at *hunchback* and *rhoNEE* enhancer-driven transcription sites to follow how the regulatory preferences of a gene changes when it is expressed at different stages of embryo development and in different locations on the embryo. In sum, we found that specific nuclear environments are associated with active transcription and that the nuclear space becomes progressively more heterogeneous as development proceeds, paralleling macroscopic morphogenesis of the whole embryo. This change in nuclear organization could alter the regulatory properties at different locations in the nucleus and times during embryogenesis, facilitating increasingly complicated regulatory patterns in later stages of embryogenesis.

## Results and Discussions

### Transcription factor distributions become more heterogeneous as development progresses

Hunchback (Hb) is a transcription factor that is active through multiple developmental stages. It is a gap gene in the early embryo, establishing the classical pattern (Figure 1A) immediately following cellularization (stage 5). Hb expression moved into neuroblasts (Figure 1B, stage 8) and eventually into the ventral nerve cord (Figure 1C, stage 14) as the embryo matured. The overall pattern localized from a broad pattern n the early embryo into individual nuclei of a specific cellular lineage. To investigate if the sub-nuclear distribution of Hb also changes during development, we imaged Hb distributions in fields of nuclei across three different developmental stages using high-resolution confocal microscopy with AiryScan and super-resolution STED microscopy. At stage 5 (Figure 1D), Hb was uniformly distributed across the entire nucleus. At stage 8 (Figure 1E), Hb began to show distinct nuclear regions of higher and lower concentrations. At stage 14 (Figure 1F), this heterogeneity has progressed further, with regions of very high Hb concentration (similar to that of stage 5) juxtaposed next to regions with almost no Hb.

**Figure 1:**
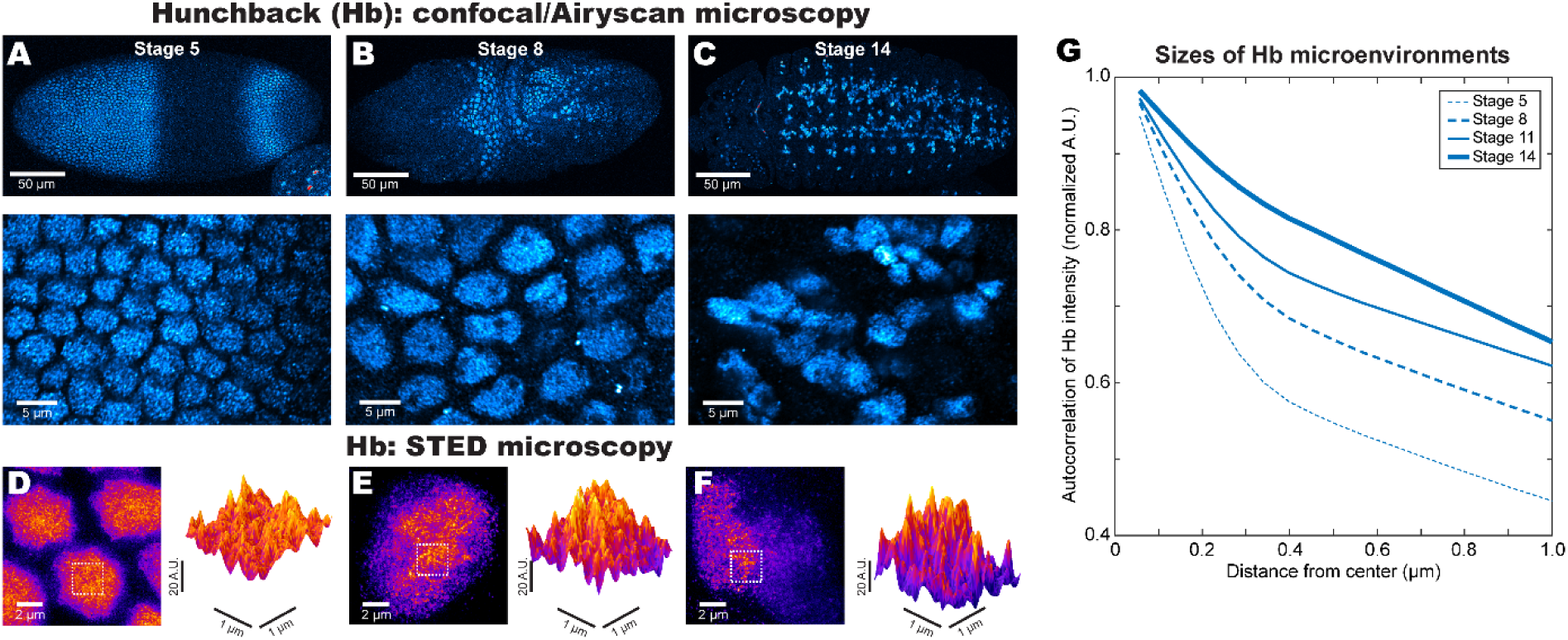
Increasing heterogeneity in Hb distribution as embryo development progresses. The distribution of Hb in *Drosophila*embryos becomes increasingly cell-specific from stage 5 (A, top panel), to stage 8 (B, top panel), and to stage 14 (C, top panel). The bottom panels in A-C show a zoomed-in view of the nuclear distribution of Hb using AiryScan. STED microscopy (D-F) shows that the levels of Hb are high and uniform across the nucleus in stage 5 (D), but becomes increasingly heterogeneous through stage 8 (E) and stage 14 (F). (G) The radially averaged autocorrelation function of Hb quantifies the overall prevalence of spatial structures at different length scales.

To quantify the spatial distributions of Hb, we computed the autocorrelation of Hb intensity (Figure 1G). The levels of correlation at a given length indicates spatial structures/features at that length scale. There was a monotonous increase in the level of correlation as the embryo ages, suggesting that localized Hb concentrations spanning several hundred nanometers became more prevalent as development proceeded. We also imaged Krüppel (Kr) distributions across similar developmental stages and computed their autocorrelation functions (Figure S1A). There was also a general upward trend, suggesting the formation of local transcription factor environments.

Other transcription factors active within a limited time frame during development, such as Ultrabithorax (Ubx, Figure S1B) and Engrailed (En, Figure S1C) showed high autocorrelation between when they are first expressed and when they reach maximum expression in the nuclei of cells in the ectoderm (stage 10 until stage 14, using STED imaging). Their autocorrelation functions decreased at stage 16, when the cells no longer expressed them and their distributions became very sparse. In sum, the nuclear distributions of Hb and Kr, which are active from early to late developmental stages, went from uniform to heterogeneous, whereas distributions of transcription factors (Ubx and En) that are only active later during development only showed the latter pattern.

### Localized histone modifications become established during development

Because transcription factors interact with the chromatin, we suspected that the state of the chromatin would influence where these factors localize. Several works have associated active regions of the chromatin with more open conformations, whereas repressed chromatin tend to be condensed (Boettiger et al., 2016; Szabo et al., 2018)(Figure S1D). We therefore imaged the distributions of several histone modifications associated using high-resolution confocal microscopy to track how their spatial distributions change as embryos age. We imaged in regions showing transcription sites from a reporter construct driven by the *hunchback* (*hb*) *cis*-regulatory region (*hbBAC*, see “Fly lines” in Materials and methods). We imaged embryos at stages 5 and 10, where nuclei transcribing from the *hbBAC* construct are within 20 μm of the surface of the embryo to preserve optimal optical resolution. We stained for H3K4me1 (Figure 2A, enhancers), H3K4me3 (Figure 2B, promoters), H3K27ac (Figure 2C, active promoters and enhancers), H3K36me3 (Figure 2D, active gene bodies) and H3K27me3 (Figure 2E, transcriptionally repressed regions).

**Figure 2:**
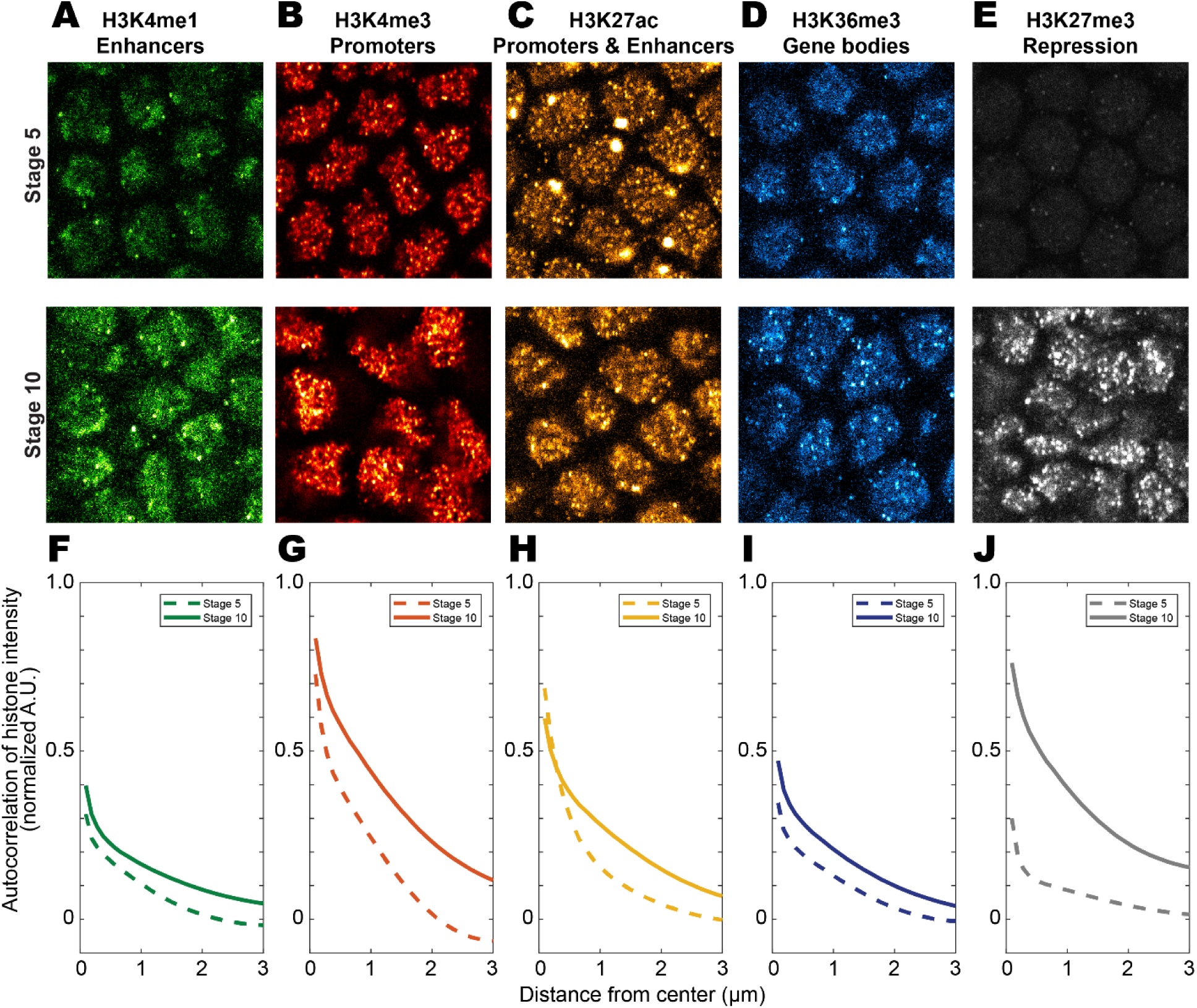
Increasing histone modifications as a function of embryo age. The chromatin shows increased levels of histone modifications as shown in (A-E) when comparing embryos at stages 5 and 10: (A) H3K4me1 marking enhancers, (B) H3K4me3 marking promoters, (C) H3K27ac marking active enhancers and promoters, (D) H3K36me3 marking active gene bodies, and (E) H3K27me3 marking repressed regions. The autocorrelation functions (F-J) of the histone modifications shows this general trend, with an increase in association across all length scales.

In general, the trend was an increasing amount of histone modifications as the embryo aged from stage 5 to 10 (Figure 2A-E). The most drastic increase was with H3K27me3, which went from barely detectible to condensed regions of very high intensities. The autocorrelation functions of the histone modifications in Figure 2F-J showed that histone marks also became more spatially expansive, with all marks showing increased correlation across all length scales. H3K27me3 (Figure 2J) showed the largest amount of change followed by H3K4me3 (Figure 2G). Overall, histone modifications became established and more heterogeneous in older embryos, but repressive modifications showed the most striking increase.

### Temporally dependent changes in the levels of histone modifications around *hunchback* transcription sites

The changing histone modification patterns during embryogenesis could alter the chromatin environments around genes active across multiple developmental stages, such as *hunchback* (*hb*). We therefore imaged how the levels of different histone modifications around *hb* transcription sites change in embryos of two different ages. To obtain strong fluorescence signals for histone modifications using immunofluorescence (IF) staining, we employed a method that does not require the denaturing step in RNA fluorescence *in situ* hybridization (RNA FISH) (Figure 3A, see “Immunofluorescence staining” in Materials and methods). In brief, we crossed the *Drosophila melanogaster* line expressing a reporter mRNA containing MS2 stem loops driven by the *cis*-regulatory region of *hb* to another line that contains MCP-GFP to label the mRNA with GFP. The mRNA transcription signal was then amplified using IF, in conjunction with staining for histone modifications (Figure 3B-E for H3K4me1 and H3K27me3). The *hb* transcription sites appeared in a radial band toward the anterior of stage 5 embryos and in neuroblasts that would become the ventral nerve cord in stage 10 embryos.

**Figure 3:**
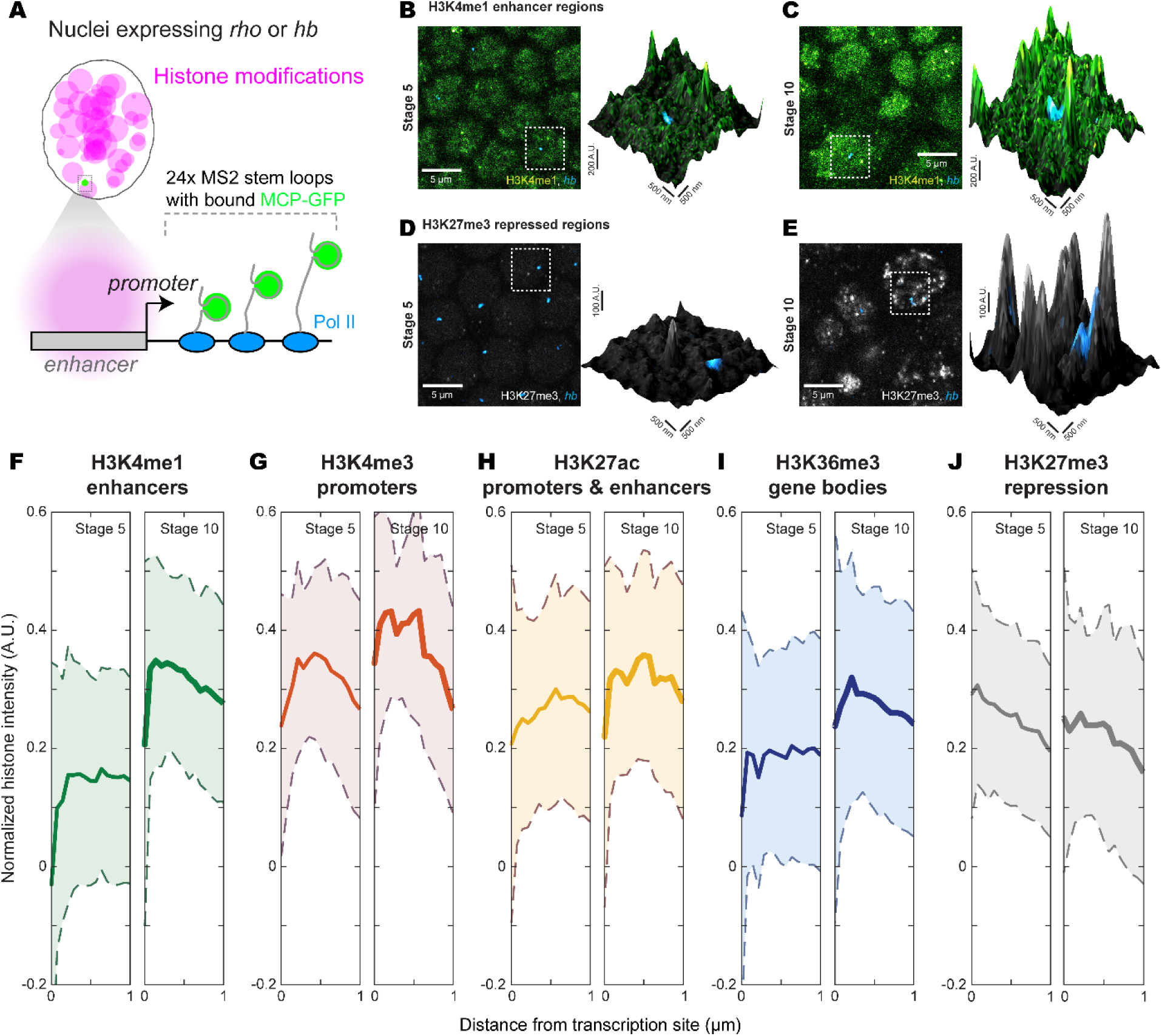
The histone environment at *hb* transcription sites depends on the timing of their expression. (A) Marking transcription sites of a MS2 reporter gene driven by the *cis*-regulatory region of *hb*or *rho*using MCP-GFP and IF amplification preserves the signal of histone modifications stained using IF. This allows the simultaneous imaging of transcription sites and histone modifications with good signal-to-noise ratio (B & C for H3K4me1 and D & E for H3K27me3, at stage 5 and 10 respectively). (F-J) The radially averaged distributions of histone modifications around *hb*transcription sites show locally specific changes between stages 5 and 10. H3K4me1: stage 5 n = 51 transcription sites, stage 10 n = 111. H3K4me3: stage 5 n = 69, stage 10 n = 52. H3K27ac: stage 5 n = 178, stage 10 n = 63. H3K36me3: stage 5 n = 54, stage 10 n = 88. H3K27me3: stage 5 n = 235, stage 10 n = 66.

To quantify the spatial distribution of histone marks around transcription sites, we computed the average radial intensity distribution function centered on the transcription site (Figure 3F-J), using the method described in (Tsai et al., 2017), also see “Image processing” in Materials and methods. The histone environment around *hb* transcription sites showed changes that deviated from the global trend presented in Figure 2. Specifically, we observed a clear increase in H3K4me1 around *hb* transcription sites going from stage 5 to stage 10 (Figure 3F), while H3K4me3 (Figure 3G) only showed a small increase. This is the opposite for their global distributions (Figure 2F & G). H3K27ac (Figure 3H) did not change between stage 5 and 10, but H3K36me3 (Figure 3I) intensity showed a moderate increase. Interestingly, many histone modifications showed an initial dip in intensity up to 200-300 nm from the transcription sites. The repressive mark H3K27me3 (Figure 3J) showed a different functional form, with no initial dip and a gentle downward slope. There were no changes between stages 5 and 10, despite the strong increase in H3K27me3 throughout the nucleus going from stage 5 to 10 (Figure 2J). The specific increase in H3K4me1 could indicate the inclusion of more distal regulatory elements in older embryos. The relatively unchanged H3K4me3 profile suggests that proximal regulatory elements are in use in both the early and late stages of development.

### Spatially dependent changes in the local histone environment at *rho* transcription sites

As we observed changes in the local histone environment as a function of time, we investigated if the local histone modification levels could also have spatial dependence on the location of the cell. The *rho* enhancer drives expression in an anterior to posterior band along the side of stage 5 embryos (Figure 4A). The sharp ventral edge of the *rho* pattern results from Snail repression and the pattern trails off gradually in the dorsal direction due to a decreasing gradient of the activator Dorsal (Figure 4B). While most transcriptionally active cells are in the active region (Figure 4C, Active), some nuclei expressing from the *rho* enhancer could still be observed in the repressed region (Figure 4C, Rep.) using a similar method for imaging *hb*transcription and histone modifications (see “Fly lines” in Materials and methods).

**Figure 4:**
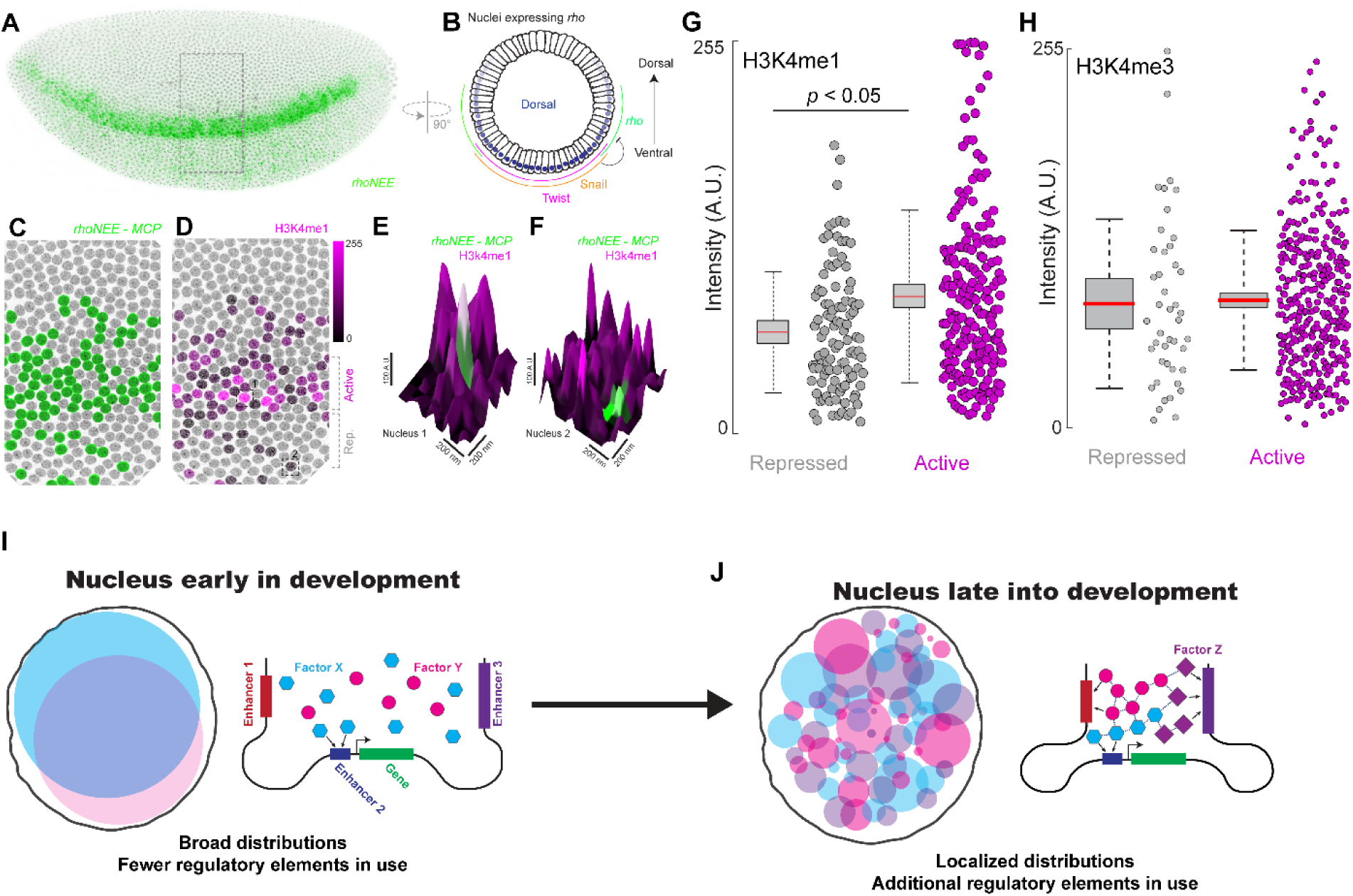
The histone environment at *rho* transcription sites depends on their location in the embryo. (A) The expression pattern of the *rhoNEE*enhancer on a side view of a stage 5 embryo. (B) The regulatory inputs of *rho* along the dorsal-ventral axis. (C) Zoomed panel from the box in (A) with nuclei with *rho* transcription site in green. (D) Same view as (C) with the shade of magenta depicting the intensity of H3K4me1 at the transcription site. (E & F) Zoomed in view around a transcription of a nucleus from the (E) active and (F) repressed zone of *rho* expression, with the z-axis showing H3K4me1 intensity. (G & H) Quantification of (G) H3K4me1 and (H) H3K4me3 levels at *rho* transcription sites. H3K4me1 repressed: n = 94 transcription sites, active: n = 535. H3K4me3 repressed: n = 44, active n = 354. (I) The nuclear environment in early embryo development is more uniform and gene regulation may preferentially occur using proximal elements. (J) The nucleus is heterogeneous in older embryos and more distal regulatory elements may be engaged in transcriptional microenvironments.

We quantified H3K4me1 and H3K4me3 at *rho* transcription sites (Figure 4D), sorting cells based on their location in the *rho* expression pattern (e.g. Figure 4E & F). We observed that transcription sites in the active region had a higher level of H3K4me1 compared to sites in the repressed region (Figure 4G, p < 0.05, two-tailed Student’s t-test). In contrast, there was no difference for H3K4me3 (Figure 4H). While the difference in H3K4me1 could be due to regulatory actions from the *rho* enhancer due to the levels of the Snail repressor, H3K4me3 levels could be explained by the reporter construct containing a poised promoter. Thus, our results show that local histone modifications could report on the state of regulatory elements, which may differ depending on the time and place of expression, even for the same gene.

### Increasing heterogeneity in the nucleus could be an integral part of embryogenesis

During development, the embryo undergoes changes in its transcriptional profiles, adapting to different regulatory needs (Figure 4I & J). In early embryos, cell division occur quickly and transcription factors form gradients spanning the embryo with uniform intra-nuclear distributions (Figure 4I). This early stage lasts approximately 6 hours in *D. melanogaster* embryos. Subsequently, the division rate slows down and the detailed segmentation of the body takes place. This phase lasts several times longer, requiring approximately 18 hours. During this phase, many regulatory and signaling pathways in different cell types and organs become active, leading to a complicated regulatory environment in the nucleus (Figure 4J). We observed that the expression patterns of transcription factors became cell-specific and their distributions within the nucleus, heterogeneous. For transcription factors that could regulate different genes during later stages of development, physical separation from early genes in the nucleus can prevent inadvertent crosstalk.

As transcription factor interacts with DNA, this partitioning of their nuclear distribution could reflect a change in the chromatin environment. Starting from when zygotic gene expression turns on, the levels of histone modifications rapidly increase in specific places on the genome. This occurs concurrent to the establishment of higher order structures, such as topologically associating domains (TADs) and to the chromatin organizing into distinct transcriptionally active or repressed regions. We observed such increases in active and repressive histone modifications in post zygotic transition embryos. The increase in H3K27me3, associated with the polycomb group proteins (Denholtz et al., 2013), was the most drastic, which could be differentiating cells shutting down genomic regions. This change in chromatin organization could create distinct accessibility patterns, guiding transcription factors into heterogeneous distributions in older embryos.

These changes in the distributions of transcription factors and chromatin states could alter the regulation of genes depending on the timing of their expression. For example, we observed an increase in H3K4me1 near active *hb* transcription sites between stage 5 and 10 embryos. In contrast, the levels of H3K4me3 were high in both younger and older embryo. The rapid division cycle and relative lack of structure in the nucleus in the early embryo might mean that genes preferentially utilize proximal regulatory elements due to kinetic constraints. As the nuclear environment becomes more stable and heterogeneous in older embryos, long-distance interactions could form and introduce distal regulatory elements. For genes active during multiple developmental stages, this permits them to respond to different regulatory inputs and serve multiple roles using the same *cis*-regulator region. Moreover, we observed a similar trend in space for the transcription sties from the *rho* enhancer, where changing H3K4me1 but constant H3K4me3 suggests a shared active promoter but differential engagement from enhancers. Together, these results imply that the local histone environments around genes could be associated with the states of different *cis*-regulatory elements. These differences could refine the expression pattern over time, as is the case for many developmental genes (Briscoe and Small, 2015), where differences in local epigenetic marking could guide incorrectly expressing or silent cells into their proper state.

This survey of the distributions of transcription factors and histone modifications in *Drosophila* embryos has provided evidence for the emergence of a heterogeneous nucleus during development. In a way, this nuclear process mirrors the increasingly complex patterns that control morphogenesis across the entire embryo. This shift in the nuclear environment highlights the need to investigate gene regulation beyond the switchover to zygotic expression and even gastrulation. Lineage-, location-, and time-specific observation of the environments around individual genes could yield insights into how regulatory mechanisms may differ between older and younger embryo. Changes in heterogeneity across different length scales could be a hallmark of embryo development and other regulatory processes in eukaryotes.

## Materials and methods

### Fly lines

All *Drosophila melanogaster* strains were maintained under standard laboratory conditions at 23 °C. *D. melanogaster* containing the *cis* -regulatory region of *hunchback* (*hb*) on a bacterial artificial chromosome (BAC) driving a *yellow* reporter mRNAs containing 24 MS2 stem loops generated in (Bothma et al., 2015) was used as a readout for the expression of *hb* . The reporter construct for *rho* is derived from the plasmid pIB-hbP2-P2P-lacZ-αTub3 UTR (Chen et al., 2012) with the hbP2 region between the restriction sites HindIII and AscI replaced with the *rhoNEE* sequence. The plasmid was synthesized by Genscript (Piscataway, New Jersey, USA). The construct was injected into Bloomington Drosophila Stock Center line 27388 and integrated using RMCE by GenetiVision (Houston, Texas, USA). Virgin flies from a transgenic *D. melanogaster* line containing MS2 coat protein (MCP) fused with eGFP driven by the *nanos* promoter from (Garcia et al., 2013) was crossed with the *hbBAC* or the *rho* reporter line to label the transcriptions sites for further amplification using immunofluorescence (IF) staining. The MCP-GFP line does not contain a nuclear localization signal (NLS) to minimize MCP-GFP forming aggregates in the nucleus in the absence of MS2 mRNAs.

### Enhancer sequence for *rhoNEE*

CTTGGGCAGGATGGAAAAATGGGAAAACATGCGGTGGGAAAAACACACATCGCGAAACATTTGGCGCAACTTGCGGAAGACAAGTGCGGCTGCAACAAAAAGTCGCGAAACGAAACTCTGGGAAGCGGAAAAAGGACACCTTGCTGTGCGGCGGGAAGCGCAAGTGGCGGGCGGAATTTCCTGATTCGCGATGCCATGAGGCACTCGCATATGTTGAGCACATGTTTTGGGGGAAATTCCCGGGCGACGGGCCAGGAATCAACGTCCTGTCCTGCGTGGGAAAAGCCCACGTCCTACCCACGCCCACTCGGTTACCTGAATTCGAGCT

### Immunofluorescence staining

Embryos from *w* ^*1118*^flies (for imaging transcription factors) and from crosses of the *hbBAC* or *rho* reporter line with the MCP-GFP line (for imaging transcription sites and histone modifications) were collected, fixed and stained using the protocols in (Crocker et al., 2015; Tsai et al., 2017). Primary antibodies were detected using secondary antibodies labeled with Alexa Fluor dyes (at a dilution ratio of 1:500, Invitrogen) for confocal and Airyscan imaging. Transcription sites, Hb and Kr were imaged using Alexa488 and the rest with Alexa555. For STED microscopy, secondary antibodies with Alexa594 (1:500, Invitrogen) or STAR RED (1:250, Abberior) were used. Ubx was imaged using Alexa594 and the rest using STAR RED.

The primary antibodies and their dilution ratios were:

Hb: Generated for the Berkeley Drosophila Transcription Network Project (Li et al., 2008), (1:100)

Kr: Generated for the Berkeley Drosophila Transcription Network Project (Li et al., 2008), (1:100)

Ubx: Developmental Studies Hybridoma Bank, FP3.38-C, (1:20)

En: Santa Cruz Biotechnology, (d-300), sc-28640, (1:50)

GFP: ThermoFisher, GFP Monoclonal Antibody (3E6), A-11120, (1:500)

H3K4me1: Merck, Anti-monomethyl-Histone H3 (Lys4) Antibody, 07-436, (1:250)

H3K4me3: Cell-Signaling Technology, Tri-Methyl-Histone H3 (Lys4) (C42D8) Rabbit mAb, 9751, (1:250)

H3K27ac: Active Motif, Histone H3K27ac antibody (pAb), 39133, (1:250)

H3K27me3: Active Motif, Histone H3K27me3 antibody (pAb), 39157, (1:250)

H3K36me3: abcam, Anti-Histone H3 (tri methyl K36) antibody, ab194677, (1:250)

### Confocal and Airyscan microscopy

Confocal and Airyscan imaging follow the protocols in (Tsai et al., 2017). Specifically, mounting of fixed Drosophila embryos was done in ProLong Gold + DAPI (Molecular Probes, Eugene, OR). Fixed embryos at the appropriate stage and orientation were imaged on a Zeiss LSM 880 confocal microscope with FastAiryscan (Carl Zeiss Microscopy, Jena, Germany). Excitation lasers with wavelengths of 405, 488 and 561 nm were used as appropriate for the specific fluorescent dyes. Whole-embryo overviews were imaged using a Zeiss LD LCI Plan-Apochromat 25x/0.8 Imm Korr DIC M27 objective. High resolution confocal and Airyscan stacks were imaged using a Zeiss Plan-Apochromat 63x/1.4 Oil DIC M27 objective. The optimal resolution as recommended by the Zen software from Zeiss was used for the x-y (70.6 nm for confocal and 42.5 nm for Airyscan) and z direction (320 nm for confocal and 190 nm for Airyscan) of stacks used for quantification. The laser power and gain were adjusted to maximize the signal to noise ratio within the dynamic range of the PMT or Airyscan detector. The acquired Airyscan stacks were processed with Zen 2.3 SP1 (Carl Zeiss Microscopy, Jena, Germany) in 3D mode to obtain super-resolved images.

### STED microscopy

Mounting of fixed Drosophila embryos was done in ProLong Diamond without DAPI (Molecular Probes, Eugene, Oregon, USA). Fixed embryos at the appropriate stage and orientation were imaged on a STEDYCON (Abberior Instruments, Göttingen, Germany) in 2D STED mode on an Olympus BX53 microscope (Olympus, Tokyo, Japan). Excitation lasers with wavelengths of 594 and 640 nm were used as appropriate for the specific fluorescent dyes. The wavelength of the STED laser was 775 nm. All samples were imaged using an Olympus UPlanSApo 100X Oil/1.4 objective. The resolutions used for the x-y and z direction were 25 nm and 250 nm, respectively. The laser power and gain were adjusted to maximize the signal to noise ratio within the dynamic range of the APD detector on the STEDYCON.

### Image processing

Image processing was done using Fiji (Schindelin et al., 2012) with the 3D ImageJ Suite plugin (Schmid et al., 2010). Image processing to obtain radially averaged intensity distributions around transcription sites was done according to (Tsai et al., 2017), with the following modification to signal normalization: for each transcription site, the maximum intensity measured in the entire radial distribution is normalized to 1, instead of normalizing to the intensity value at *r* = 0. Radially averaged autocerrelation functions were computed using an ImageJ macro (imagejdocu.tudor.lu/macro/radially_averaged_autocorrelation). AiryScan images were used for their wider field of view compared to STED, in order to include a larger number of nuclei. Intensity distributions and autocorrelation fuctions were plotted using Matlab (MathWorks, Natick, MA).

## Acknowledgements

The *hunchback* BAC line was a generous gift from Michael Levine. The MCP-eGFP line and the plasmid pIB-hbP2-P2P-lacZ-αTub3 UTR were generous gifts from Hernan Garcia. The Hunchback and Krüppel primary antibody were generous gifts from Mark Biggin. We would like to thank Marko Lampe at the EMBL Advanced Light Microscopy Facility for his expert advice and training to conduct STED imaging. We would also like to thank members of the Crocker Group for valuable discussions and suggestions to improve the manuscript.

## Data and Material Availability

The original images (AiryScan, confocal, and STED) will be available for download and are indexed at: https://www.embl.de/download/crocker/XXXXX. All fly lines and plasmid sequences will be available upon reasonable request.

---

**Figure S1:**
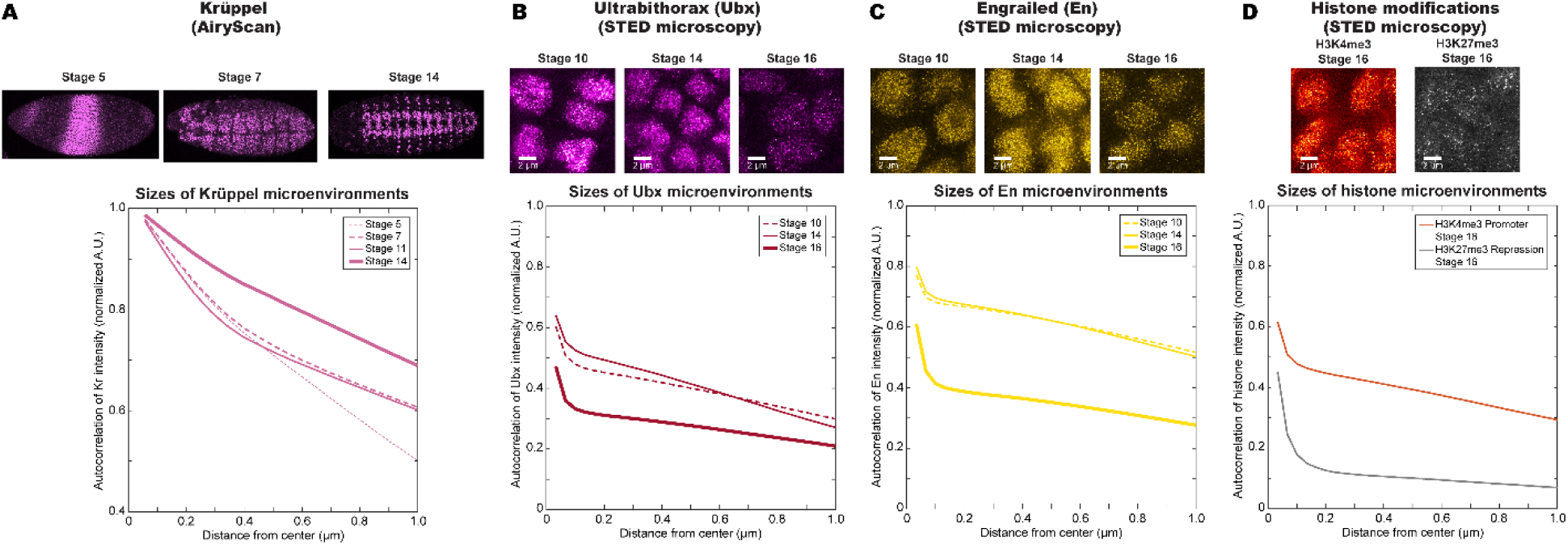
The spatial scales of transcription factors and histone marks as a function of embryo age and regulatory function. (A) An increase in the autocorrelation function is also observed for Krüppel across developmental stages. Transcription factors that are only expressed during later stages of embryo development, such as (B) Ubx and (C) En, do not show a uniform nuclear distribution. During stages 10 and 14, when both factors are expressed, their distributions are heterogeneous and show significant levels of autocorrelation. Their expression decreases significantly by stage 16. (D) The physical sizes of the chromatin can vary according to their transcriptional state.

